# Voltage Gated Calcium Channel Dysregulation May Contribute to Neurological Symptoms in Calmodulinopathies

**DOI:** 10.1101/2024.12.02.626503

**Authors:** John W. Hussey, Emily DeMarco, Deborah DiSilvestre, Malene Brohus, Ana-Octavia Busuioc, Emil D. Iversen, Helene H. Jensen, Mette Nyegaard, Michael T. Overgaard, Manu Ben-Johny, Ivy E. Dick

**Author notes:** Address correspondence to: Ivy E. Dick, University of Maryland, School of Medicine, Baltimore, MD 21210.

## Abstract

Calmodulinopathies are caused by mutations in calmodulin (CaM), and result in debilitating cardiac arrythmias such as long-QT syndrome (LQTS) and catecholaminergic polymorphic ventricular tachycardia (CPVT). In addition, many patients exhibit neurological comorbidities, including developmental delay and autism spectrum disorder. Until now, most work into these mutations has focused on cardiac effects, identifying impairment of Ca^2+^/CaM-dependent inactivation (CDI) of Ca_V_1.2 channels as a major pathogenic mechanism. However, the impact of these mutations on neurological function has yet to be fully explored. CaM regulation of voltage-gated calcium channels (VGCCs) is a critical element of neuronal function, implicating multiple VGCC subtypes in the neurological pathogenesis of calmodulinopathies. Here, we explore the potential for pathological CaM variants to impair the Ca^2+^/CaM-dependent regulation of Ca_V_1.3 and Ca_V_2.1, both essential for neuronal function. We find that mutations in CaM can impair the CDI of Ca_V_1.3 and reduce the Ca^2+^-dependent facilitation (CDF) of Ca_V_2.1 channels. We find that mutations associated with significant neurological symptoms exhibit marked effects on Ca_V_1.3 CDI, with overlapping but distinct impacts on Ca_V_2.1 CDF. Moreover, while the majority of CaM variants demonstrated the ability to bind the IQ region of each channel, distinct differences were noted between Ca_V_1.3 and Ca_V_2.1, demonstrating distinct CaM interactions across the two channel subtypes. Further, C-domain CaM variants display a reduced ability to sense Ca^2+^ when in complex with the Ca_V_ IQ domains, explaining the Ca^2+^/CaM regulation deficits. Overall, these results support the possibility that disrupted Ca^2+^/CaM regulation of VGCCs may contribute to neurological pathogenesis of calmodulinopathies.

## Introduction

Calmodulin (CaM) is a small yet indispensable protein encoded by the *CALM1, CALM2,* and *CALM3* genes that is found in all eukaryotic cells where it acts as an essential Ca^2+^ sensor (Hussey et al., 2023). Changes in cytosolic Ca^2+^ levels encode information that is critical for cellular function, and CaM is readily able to detect these changes via Ca^2+^ binding at four EF hand domains. These EF hands occur in coordinating pairs, with two in each of the two lobes of CaM (**Fig. 1A**). When CaM binds Ca^2+^, it engages in a rapid conformational change, imparting Ca^2+^ dependent effects to an array of target proteins (Westerlund and Delemotte, 2018), including voltage gated calcium channels (VGCCs) (Ben-Johny and Dick, 2022). Interestingly, each lobe of CaM is capable of imparting distinct and independent forms of regulation on target proteins (DeMaria et al., 2001; Ben-Johny and Dick, 2022). Thus, CaM provides a vital function in diverse cellular processes that range from cellular differentiation and gene expression to muscle contraction and membrane excitability.

**Figure 1:**
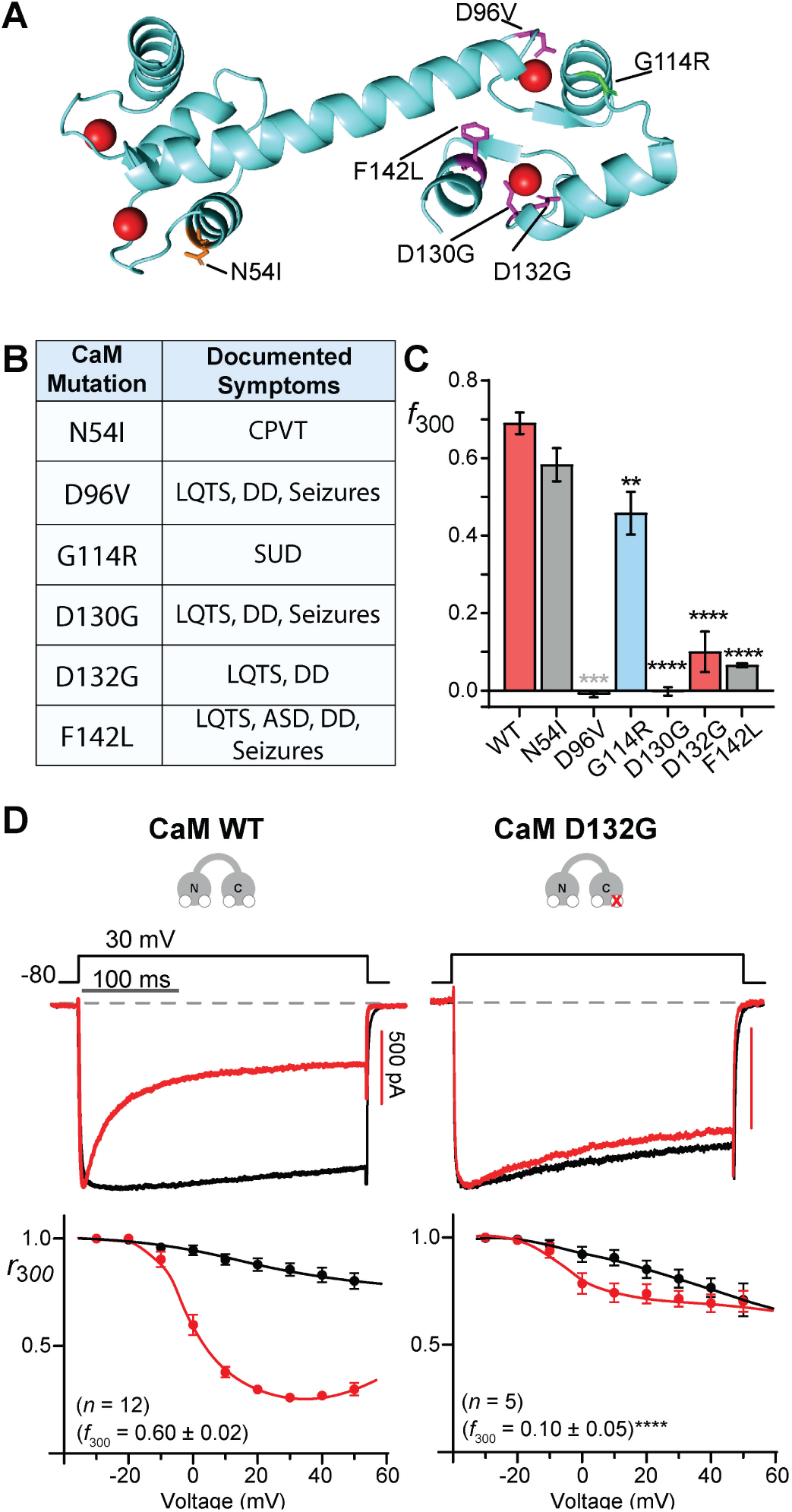
CaM mutants impair Ca_V_1.2 CDI. **A**: Depiction of CaM mutation described in this study; primarily associated with CPVT (orange), LQTS and neurodevelopmental delay (pink), and sudden death (SUD) (green) highlighted. **B**. Table describing the reported clinical phenotypes associated with each CaM mutation. Symptoms were originally reported in (Nyegaard et al., 2012; Crotti et al., 2013; Kaplanis et al., 2020; Crotti et al., 2023) **C**: Degree of Ca_V_1.2 CDI conferred by CaM mutants. CDI is evaluated as the difference in the fractional Ba^2+^ and Ca^2+^ currents remaining after 300 ms following sustained depolarizations to 30 mV from -80 mV. Data in gray (Limpitikul et al., 2014), and blue (Brohus et al., 2021) were previously reported. **D**, Top: exemplar traces displaying inactivation of Ca_V_1.2 currents in presence of CaM WT or CaM D132G, with Ba^2+^ currents (black) normalized to Ca^2+^ currents (red). Scale bars reflect Ca^2+^ currents. Bottom: data displaying *r*_300_ values at varying test potentials. CDI evaluated *f*_300_ at 30 mV was compared via a Student’s t-test (**** *p* < 0.0001). Data plotted ± SEM.

Over the past decade, a growing number of missense mutations in CaM have been identified in patients suffering from debilitating and life-threatening cardiac arrhythmias, known as calmodulinopathies (Nyegaard et al., 2012; Crotti et al., 2013; Limpitikul et al., 2014; George, 2015; Reed et al., 2015; Crotti et al., 2019; Nyegaard and Overgaard, 2019; Crotti et al., 2023; Hussey et al., 2023). Disease phenotypes primarily include long-QT syndrome (LQTS) and/or catecholaminergic polymorphic tachycardia (CPVT) (**Fig. 1A**)(Nyegaard et al., 2012; Crotti et al., 2023). Investigations revealed multiple pathological mechanisms attributable to the calmodulinopathy variants, typically involving diminished ability of CaM to bind to Ca^2+^ (Limpitikul et al., 2014). Notably, almost all LQTS-associated variants identified thus-far are located within the C-lobe of CaM (Crotti et al., 2023; Hussey et al., 2023). Many of these variants were found to impair the Ca^2+^/CaM-dependent regulation of the VGCC Ca_V_1.2 (Limpitikul et al., 2014; Limpitikul et al., 2017), an essential component of cardiac function. In these channels, CaM dependent regulation takes the form of Ca^2+^/CaM-dependent inactivation (CDI), a vital form of negative feedback that tempers Ca^2+^ entry into cells. While each lobe of CaM is independently capable of imparting CDI, the majority of Ca_V_1.2 CDI stems from the C-lobe, rationalizing the importance of this locus in the context of calmodulinopathies. Disruption of CDI in Ca_V_1.2 is associated with Timothy syndrome (TS), a multisystem disorder often presenting with symptoms similar to that of calmodulinopathies including LQTS, neurodevelopmental delay and seizures (Timothy et al., 2024).

Ca_V_1.2 channels are thought to be particularly vulnerable to CaM mutations because of the manner in which the two proteins interact. Many targets of CaM regulation bind preferentially to the calcified form of the protein; Ca_V_1.2 channels, by contrast, preassociate with Ca^2+^-free CaM (apoCaM) via the IQ motif within the carboxy-tail of the channel (Lee et al., 1999; DeMaria et al., 2001; Erickson et al., 2001). In the case of non-preassociated targets, a loss of Ca^2+^ binding within a small subset of cellular CaM proteins (as occurs in calmodulinopathies) has minimal impact in the presence of sufficient WT CaM. However, when CaM is preassociated with its target in the apo state, some fraction of target proteins will be bound to a mutant CaM, and are thus unable to undergo Ca^2+^ dependent regulation (Hussey et al., 2023). In the case of calmodulinopathies a heterozygous mutation within a single *CALM* allele results in only one effected allele out of six (Schwartz et al., 2024), yet this limited expression of the mutant has been shown to be sufficient to diminish CDI in the context of a ventricular myocyte (Limpitikul et al., 2014). In this way, loss of CDI in Ca_V_1.2 due to mutations in CaM contributes strongly to the LQTS phenotype (Limpitikul et al., 2014).

Though the primary reported clinical finding in calmodulinopathies has been cardiac in nature, a number of cases present with additional neurological comorbidities, including developmental delay, seizures, and autism spectrum disorder (Crotti et al., 2023). While initial reports characterized consequences of CaM mutations as leading to LQTS or CPVT, more recent work highlights intermediate or mixed phenotypes; moreover, some CaM mutations have been identified in patients with predominantly neurological symptoms and minimal or no identified cardiac pathology (Crotti et al., 2023). Further, recent work in *Caenorhabditis elegans* has shown that calmodulinopathy mutations can directly impair neuronal function (Jensen et al., 2023). To what extent, and by which mechanism, these CaM variants may be contributing to the observed neurological symptoms in patients has yet to be elucidated. Notably, Ca_V_1.2 is highly expressed in the brain (Tang et al., 2004), and neurological symptoms represent a common feature of mutations in this channel (Herold et al., 2023). Moreover, the brain is known to express multiple CaM regulated Ca_V_ channel subtypes, implicating these channels in the neurological pathogenesis of calmodulinopathies.

The features that make Ca_V_1.2 particularly vulnerable to calmodulinopathy mutations are shared by other Ca_V_ channel subtypes. In particular, Ca_V_1.3 and Ca_V_2.1 have broad structural similarity to Ca_V_1.2, harbor similar CaM binding IQ motifs, exhibit robust C-lobe CaM mediated regulation, and demonstrate preassociation (Lee et al., 1999; Erickson et al., 2001; Ben-Johny and Dick, 2022). Moreover, these two channels are widely expressed across multiple brain regions (Hell et al., 1993; Tan et al., 2011), contribute to long-term potentiation (Gamelli et al., 2011), and disruption of these channels has been associated with autism-spectrum disorder (Pinggera et al., 2015), developmental delay (Folacci et al., 2023), and other neurological disorders (Pietrobon, 2010; Folacci et al., 2023). As in Ca_V_1.2, CaM modulates Ca_V_1.3 by conferring CDI, with distinct contributions from each lobe of CaM (DeMaria et al., 2001). Additionally, multiple splice variants of Ca_V_1.3 exist with distinct CaM regulatory features. While a short form of the channel (Ca_V_1.3_43s_) exhibits robust CDI, a long splice variant Ca_V_1.3_L_, contains a C-terminal domain, termed inhibitor of CDI (ICDI) (Wahl-Schott et al., 2006) or C-terminal modulatory domain (CTM) (Lieb et al., 2012), that diminishes CDI by competing with CaM binding to the channel (Liu et al., 2010). Interestingly, this splice variant exhibits increased voltage-dependent inactivation (VDI), even as CDI is reduced (Singh et al., 2008). These splice variants are crucial for adjusting the channel’s activity for distinct cellular requirements, demonstrating the importance of fine-tuned CaM regulation in physiological function (Bock et al., 2011; Ben-Johny and Dick, 2022). For Ca_V_2.1, CDI is primarily mediated by the N-lobe of CaM, while the C-lobe drives Ca^2+^-dependent facilitation (CDF), characterized by a transient elevation of whole-cell currents following Ca^2+^ entry through the channel (DeMaria et al., 2001). CDF permits rapid and dynamic modulation of Ca_V_2.1 current within the presynaptic terminal, and it is thought to contribute to short-term synaptic plasticity and rhythmicity (Catterall et al., 2013; Nanou et al., 2018).

In this study, we examine the biophysical properties of calmodulinopathy variants in the context of Ca_V_1.3 and Ca_V_2.1, both in terms of channel regulation and binding affinity. We focus on mutations identified within the C-lobe of CaM previously shown to decrease the affinity of Ca^2+^ binding to CaM and reduce the CDI of Ca_V_1.2 channels (Crotti et al., 2013). We find that many of these mutations exhibit profound disruptions of Ca_V_1.3 CDI and Ca_V_2.1 CDF, demonstrating a potential role for these channels in the neurological effects observed in calmodulinopathy patients. Comparatively, a CPVT-associated mutation within the N-lobe of CaM demonstrated no significant impact on CDI or CDF of these channels, consistent with a primarily cardiac pathology acting through the ryanodine receptor (RyR2), as previously described (Hwang et al., 2014; Sondergaard et al., 2015b; Sondergaard et al., 2020). Thus, our results demonstrate a potential mechanism by which LQTS-associated CaM variants may also influence neurological health through their impact on multiple VGCCs in the brain.

## Results

### Ca_V_1.2 and CaM D132G

Prior work has examined the impact of calmodulinopathy variants associated with a range of symptoms. Here, we focus on several variants which have been strongly implicated in LQTS, CPVT, and sudden death, with variable reports of neurological features (**Fig. 1 A, B**). In the case of LQTS, CaM mutants are thought to contribute to the disease largely through the disruption of Ca_V_1.2 CDI (Limpitikul et al., 2014). The majority of C-lobe mutations focused on in this study have previously been shown to blunt Ca_V_1.2 CDI, while the CPVT-associated N54I mutation, located in the N-lobe, largely spares CDI (**Fig. 1C**) (Limpitikul et al., 2014; Brohus et al., 2021). However, the effect of the CaM mutation D132G has yet to be evaluated on Ca_V_1.2 despite association with severe LQTS (Zahavich et al., 2018) and developmental delay (Kaplanis et al., 2020). We therefore sought to determine the impact of CaM D132G on Ca_V_1.2 CDI. To do so, we coexpressed mutant CaM with Ca_V_1.2 in HEK293 cells, such that CDI can be seen as the larger decay of the Ca^2+^ current (**red**), as compared to the Ba^2+^ current (**black**) in response to a sustained depolarization from –80 mV to 30 mV (**Fig. 1D**, top). CDI is then quantified by comparing the ratio of Ca^2+^ current that remains following 300 ms of depolarization (*r*_300_) with that of Ba^2+^. In this way, Ba^2+^ currents serve to control for the voltage-dependent inactivation (VDI) of the channels, which occurs alongside CDI and is minimized in our recording conditions by the use of the β_2a_ auxiliary subunit. In the presence of CaM D132G, the Ca^2+^ current is largely sustained following the depolarizing step, demonstrating a loss of CDI which persists across a range of test potentials (**Fig. 1D**, bottom). Thus, CaM D132G displays a similar capacity to reduce Ca_V_1.2 CDI as compared to other previously documented LQTS-associated CaM variants (Limpitikul et al., 2014) (**Fig. 1C**).

### The Impact of Pathological CaM mutations on Ca_V_1.3 Regulation

Given that modulation of channel gating by CaM extends beyond Ca_V_1.2, we investigated the effects of pathological CaM variants on Ca_V_1.3 inactivation, starting with the Ca_V_1.3_L_ splice variant. Ca_V_1.3_L_ channels represent a long splice variant (also called Ca_V_1.3_42_ or Ca_V_1.3_8a_ (Bock et al., 2011)), which includes the ICDI/CTM (Singh et al., 2008; Lieb et al., 2012) resulting in a channel with attenuated CDI. Nonetheless, under conditions where exogenous CaM is overexpressed, robust CDI is observed in the presence of CaM WT (**Fig. 2A**). Strong CDI (**red**) is observed across a range of depolarizing potentials (**Fig. 2A**, bottom), while Ba^2+^ currents (**black**) exhibit only modest VDI. While the N-lobe CaM N54I produced no significant deficits in CDI (**Fig 2B**), each of the C-lobe CaM mutations imparted significant reduction in CDI as compared to CaM WT (**Fig. 2C-G**), although it should be noted that the CDI observed in the presence of G114R displayed more rapid inactivation kinetics as compared to the other mutants. As CaM is capable of mediating CDI in Ca_V_1.3_L_ channels through both the C- and N-lobes (Dick et al., 2008), the remaining CDI observed in the calmodulinopathy variants is likely due largely to N-lobe CDI, which is known to exhibit slower inactivation kinetics as compared to C-lobe mediated CDI (Peterson et al., 1999; Yang et al., 2006; Dick et al., 2008). Consistent with this, an artificially mutated CaM_1234_, in which Ca^2+^ binding is eliminated in both lobes of CaM results in complete abolition of CDI in Ca_V_1.3_L_ (**Fig. 2H**). Thus, calmodulinopathy variants can impair Ca_V_1.3_L_ CDI in a manner similar to that observed in Ca_V_1.2 (**Fig. 1**).

**Figure 2:**
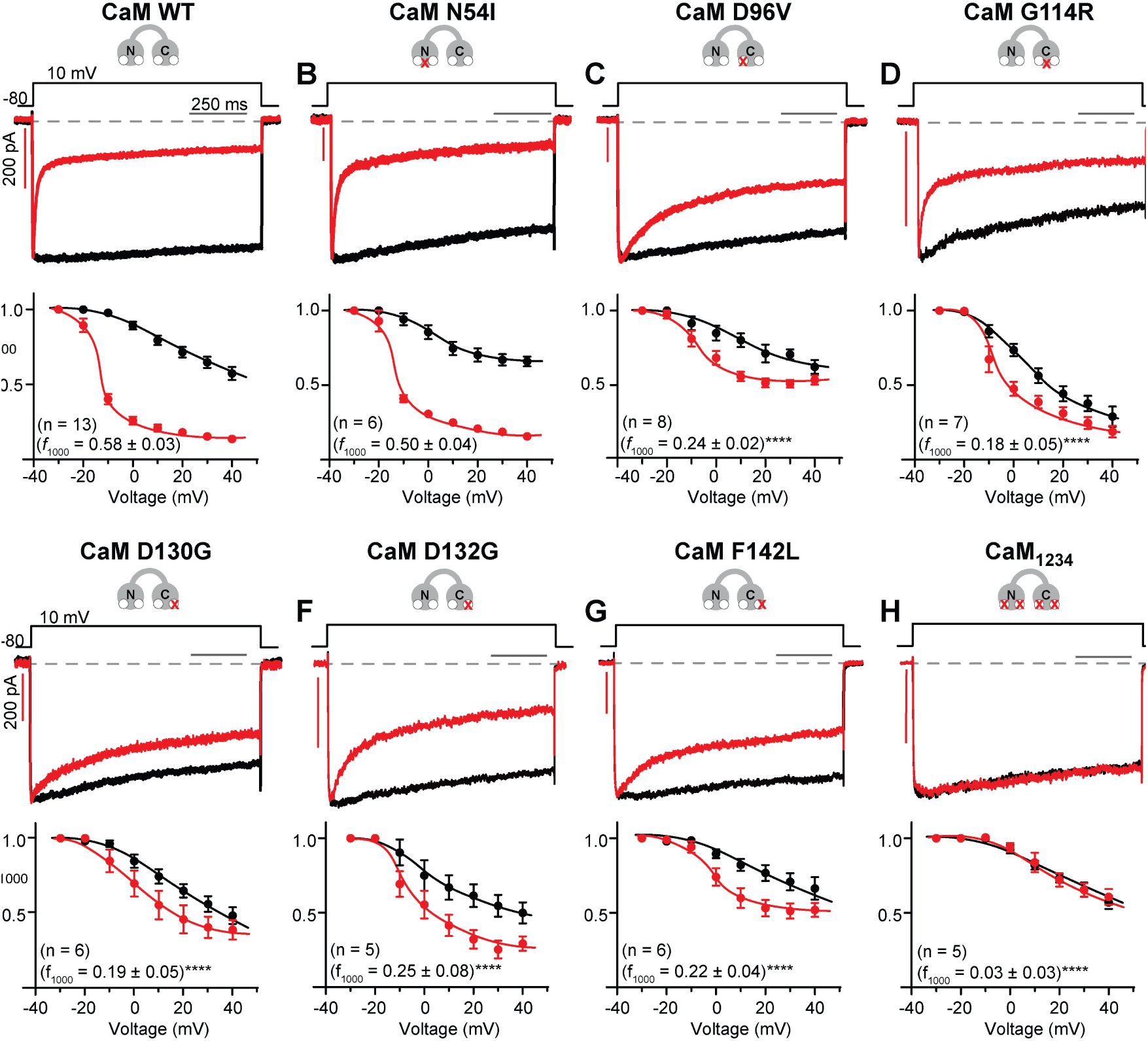
CaM mutants impair Ca_V_1.3_L_ CDI. **A-H**, Top: Exemplar current traces for Ca_V_1.3_L_ co-transfected with the indicated CaM variant. Ba^2+^ currents (black) are normalized to Ca^2+^ currents (red). Scale bars reflect Ca^2+^ currents. Bottom: population data displaying *r*_1000_ values for Ca_V_1.3_L_ and the indicated CaM variants. *f*_1000_ values were compared at 10 mV for each variant using a one-way ANOVA, followed by Dunnet’s test for multiple comparisons relative to WT (**** *p* < 0.0001). Data plotted ± SEM.

Analysis of Ba^2+^ current decay during the depolarization revealed a slight but significant variability in VDI. Specifically, at higher test potential voltages, CaM G114R imparted greater VDI than other mutants or WT CaM as revealed by Ba^2+^ *r*_1000_ values (**Fig 2D**, black). Additionally, CaM D132G appeared to follow a similar trend, but this effect did not rise to the level of significance (**Fig. 2F**). To investigate this phenomenon further, these mutants were coexpressed with Ca_V_1.3_L_ in conditions optimized to reveal voltage-dependent effects. Employing the β_1b_ accessory subunit in the transfection permits robust VDI in Ca_V_1.3_L_ channels (**Fig. 3A**, top). Once again, the G114R mutant revealed a slight but significantly enhanced VDI as compared to CaM WT or CaM D132G, as revealed by smaller *r*_1000_ values following 20 mV depolarizations (**Fig. 3A-C**, bottom panel). Thus, it appears that VDI is uniquely different in the CaM G114R recordings.

**Figure 3:**
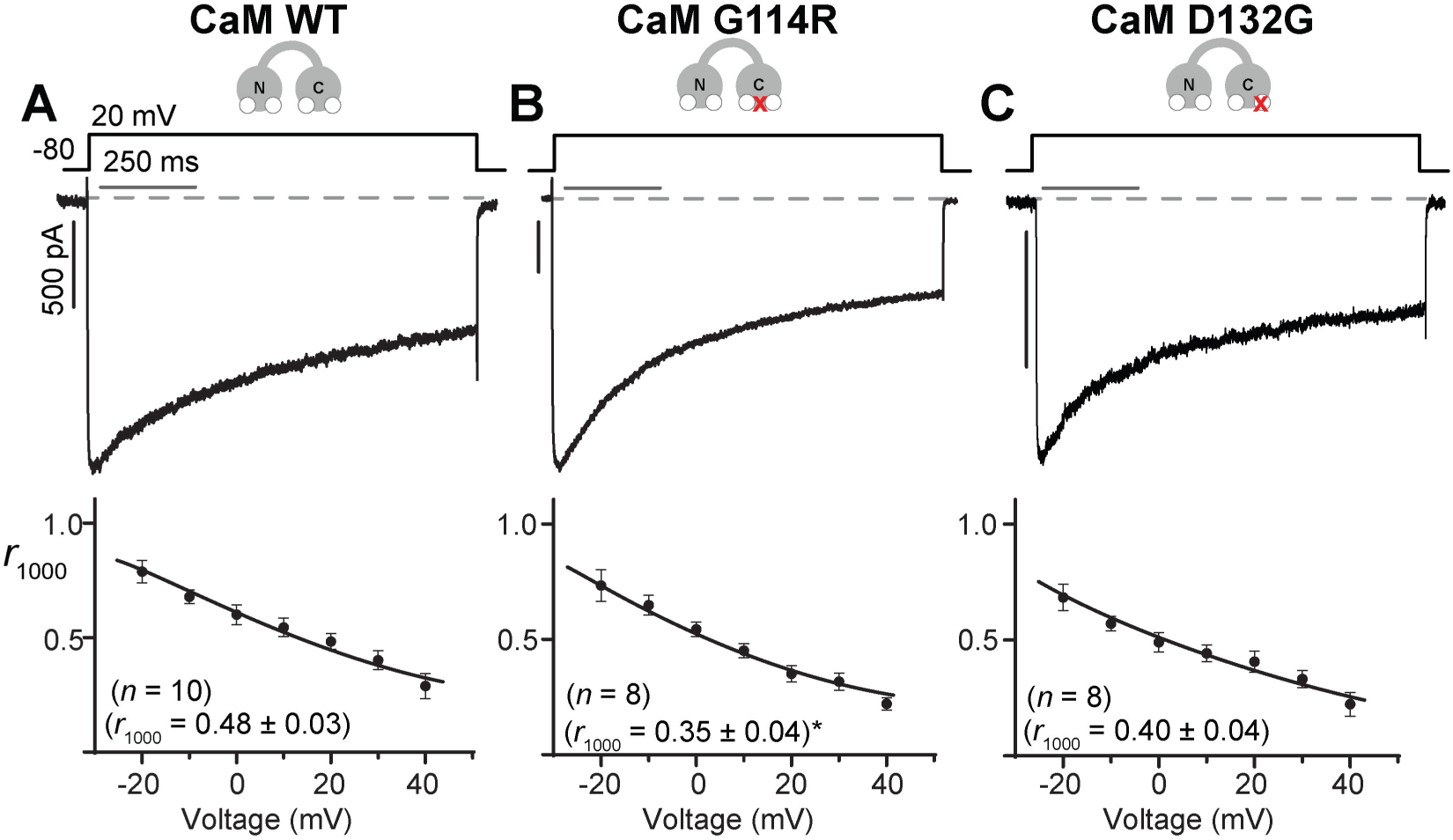
CaM variant G114R increases Ca_V_1.3_L_ VDI. **A-C,** Top: Exemplar Ba^2+^ current traces for Ca_V_1.3_L_ co-transfected with the indicated CaM variant and β_1b_ accessory subunit. Bottom: Ba^2+^ *r*_1000_ values displayed for varying test potential voltages. *r*_1000_ values were compared at 20 mV for each CaM variant via a one-way ANOVA, followed by Dunnet’s test for multiple comparisons relative to WT (* *p* = 0.035). Data plotted ± SEM.

We next considered the impact of the CaM mutants on a short isoform of Ca_V_1.3, Ca_V_1.3_43s_, which lacks the modulatory ICDI/CTM domain and thus displays full CDI (Bock et al., 2011). The CaM mutants had generally similar impacts on CDI in Ca_V_1.3_43s_ as they did in Ca_V_1.3_L_ (**Fig. 4A-H**). As expected, the N54I N-lobe CPVT mutant was not statistically different from WT (**Fig. 4B**), while mutations D96V, D130G, D132G, and F142L all significantly impaired CDI (**Fig. 4B, 4C, 4E-G**). The artificial mutation CaM_1234_ produced a near total lack of Ca_V_1.3_43s_ CDI (**Fig. 4H**). In contrast, CaM G114R, which displayed diminished CDI as compared to overexpressed CaM WT in Ca_V_1.3_L_, showed no effect on Ca_V_1.3_43s_ CDI (**Fig. 4D**). Additionally, no effects on Ca_V_1.3_43s_ VDI were observed.

**Figure 4:**
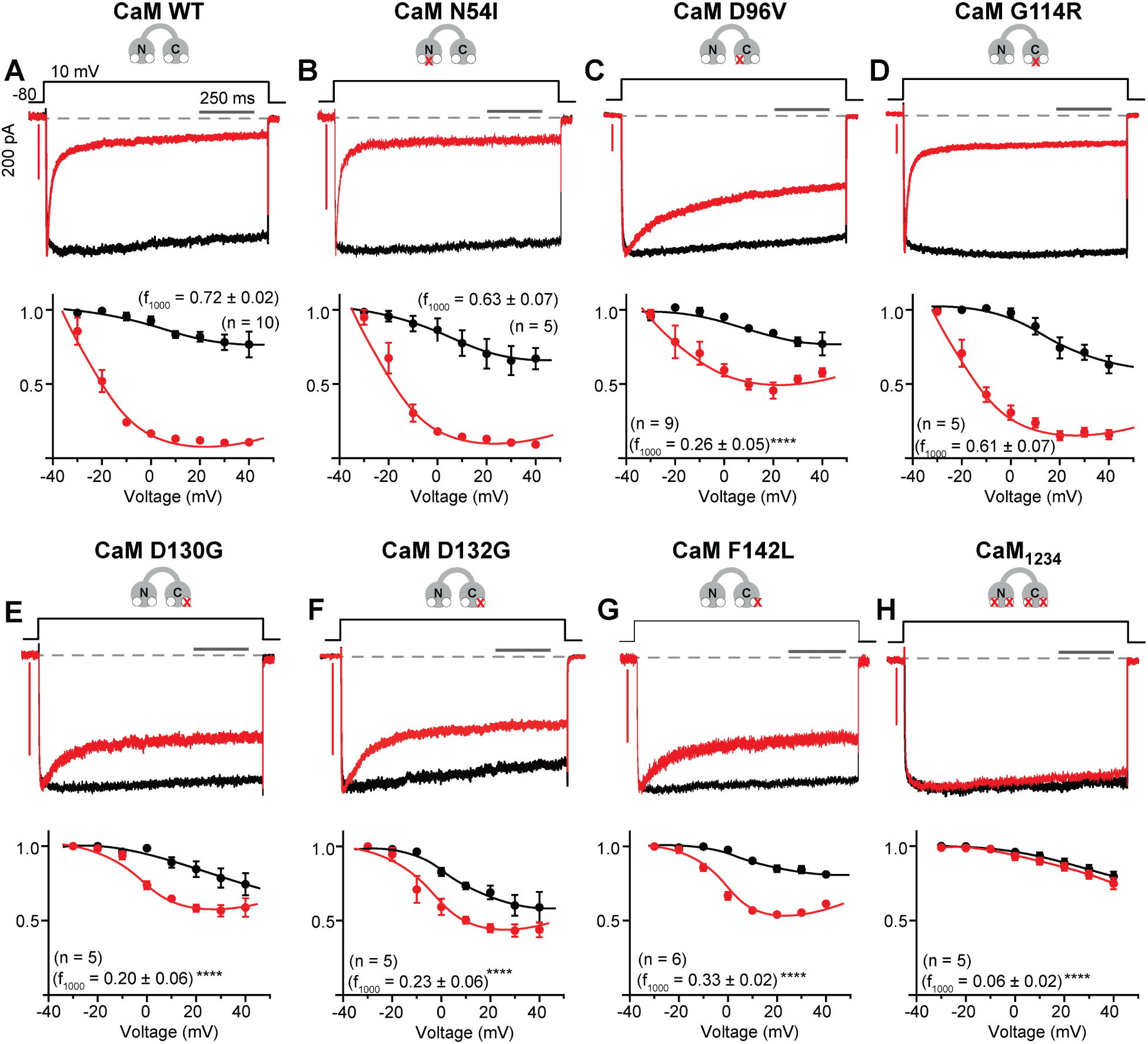
CaM mutants impair Ca_V_1.3_43s_ CDI. **A-H**, Top: Exemplar traces for Ca_V_1.3_43s_ co-transfected with the indicated CaM variant. Ca^2+^ currents (red) are normalized to Ba^2+^ currents (black), with scale bars referring to Ca^2+^. Bottom: data displaying *r*_1000_ values across varying test potentials. *f*_1000_ values were compared at 10 mV for each of the CaM variants using a one-way ANOVA, followed by Dunnet’s test for multiple comparisons as compared to WT (**** *p* < 0.0001). Data plotted ± SEM.

### The effect of CaM mutations on Ca_V_1.3 association

While most C-lobe CaM mutants conferred Ca_V_1.3_L_ and Ca_V_1.3_43s_ CDI deficits in a similar fashion to their action on Ca_V_1.2, data from CaM G114R diverges from this paradigm. It is therefore reasonable to question how CaM G114R interacts with Ca_V_1.3. Notably, the results for each Ca_V_1.3 splice variant would be consistent with a lack of CaM G114R binding to the channel. Thus, the possibility emerges that CaM G114R does not interact with either Ca_V_1.3 variant, and the results demonstrate the expected CDI under endogenous CaM WT levels (Liu et al., 2010). Moreover, any mutant CaM must bind to the channel at least as well as CaM WT in order for preassociation to exert its full impact in the context of a heterozygous mutation in one of the three *CALM* genes (Erickson et al., 2003; Limpitikul et al., 2014; Badone et al., 2018; Hussey et al., 2023). We therefore sought to explore whether each mutant CaM is capable of binding the channel with a similar affinity as compared to CaM WT utilizing Förster resonance energy transfer (FRET). To quantify binding activity, we employed a previously validated (Ben-Johny et al., 2016; Lee et al., 2016; Rivas et al., 2021) live-cell flow cytometric methodology to construct FRET-based binding curves and estimate the effective binding affinity (*K*_D,Eff_) for each channel/CaM pair. A peptide spanning known C-terminal CaM binding regions of Ca_V_1.3 was ligated to a donor fluorophore, while CaM was joined to an acceptor fluorophore (**Fig. 5A**), and the two were expressed in HEK293 cells under basal Ca^2+^ conditions. Exemplar binding curves (**Fig. 5D-I**) of CaM variants N54I, D96V, D130G, D132G, and F142L show broad similarity with the binding curve of CaM WT, and the average results from each of these mutants demonstrate no significant alterations to *K*_D,Eff_ relative to WT (**Fig. 5B**). However, CaM G114R demonstrates a profoundly diminished capacity to bind to the channel fragment (**Fig. 5B, F**), indicating reduced preassociation with the channel in the apo state.

**Figure 5:**
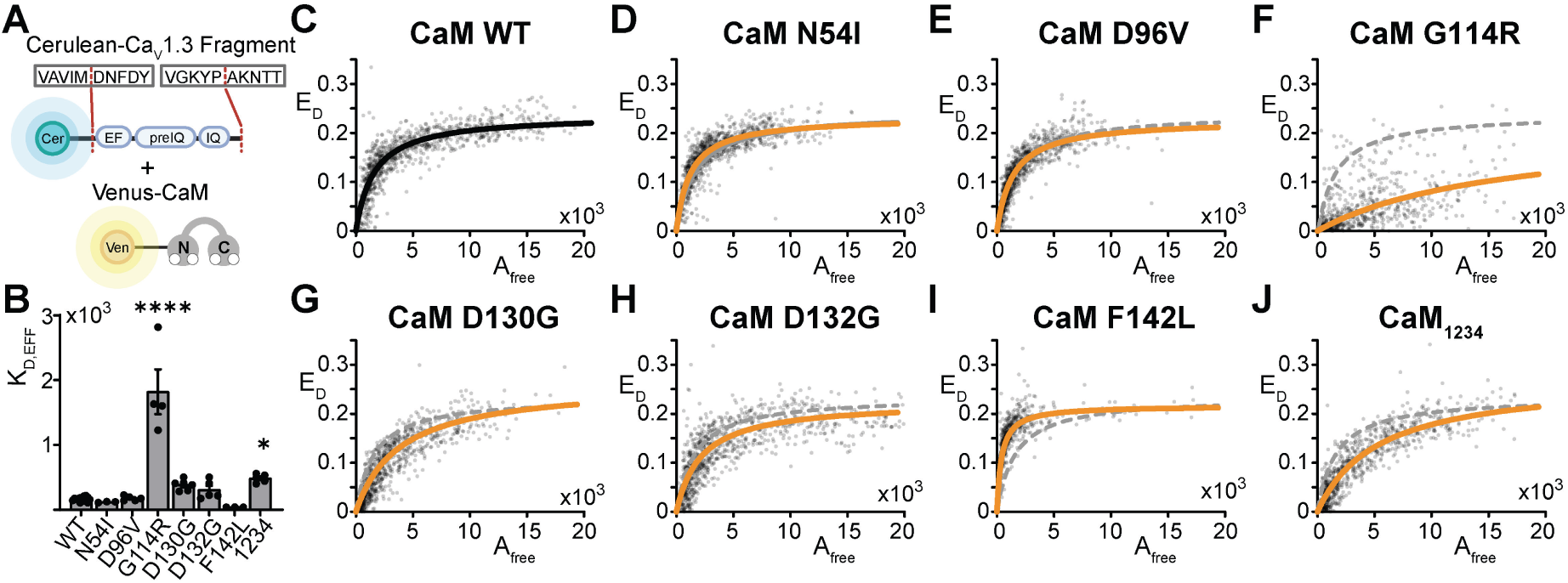
Evaluation of CaM binding to Ca_V_1.3. **A:** Illustration of the constructs utilized. A Ca_V_1.3 C-terminal channel fragment including the IQ domain as indicated is concatenated with Cerulean. Each CaM variant was generated within a Venus-CaM construct. **B**: Measured relative binding efficiencies (*K*_D,eff_) plotted for the Ca_V_1.3 channel fragment and CaM variants in low cytosolic Ca^2+^. Significance was ascertained with a one-way ANOVA followed by a Dunnet’s test for multiple comparisons relative to WT (****, *p* > 0.0001; *, *p* = 0.033). Data plotted ± SEM. **C-J**: FRET binding curves from a live-cell flow cytometric FRET assay for the Ca_V_1.3 channel fragment and indicated CaM variant. Each point reflects the result from an individual cell, with a curve in black (WT) or orange (mutant) fitted to estimate effective *K*_D_ (*K*_D,eff_). Data is plotted as a function of free acceptor fluorophore molecules (*A_f_*_ree_) and donor-centric FRET efficiency (*E*_D_). WT results are reproduced on variant data as the gray dashed curve for comparison.

Given that CaM binding can be impacted by its association with Ca^2+^ (Brohus et al., 2021), we investigated whether CaM G114R/Ca_V_1.3 binding could be recovered under conditions sufficient to calcify CaM. Subsequent testing revealed that, in the presence of an ionophore and high external Ca^2+^, binding activity could be partially rescued (**Fig. 6A-C**). These data indicate that calcified CaM G114R maintains the ability to bind Ca_V_1.3 at high Ca^2+^, but preassociation is likely reduced. Given this result, the modified CDI observed in Ca_V_1.3_L_ (**Fig. 2D)** as compared to overexpressed CaM WT **(Fig. 2A)** is likely due to the known competition between CaM and ICDI/CTM for binding to the Ca_V_1.3 IQ (**Fig. 6D**). At low, endogenous levels of CaM WT, ICDI/CTM is able to out-compete CaM for binding to the channel, resulting in a significant reduction of CDI as previously described (Wahl-Schott et al., 2006; Liu et al., 2010; Lieb et al., 2012). Upon overexpression of CaM WT, CDI is restored due to mass action (**Fig. 2A**). Without sufficient apoCaM binding, CaM G114R does not alter this scheme. Moreover, in Ca_V_1.3_43s_ lacking ICDI/CTM, endogenous CaM is sufficient to drive robust CDI, and the lack of CaM G114R binding causes no apparent difference in CDI.

**Figure 6:**
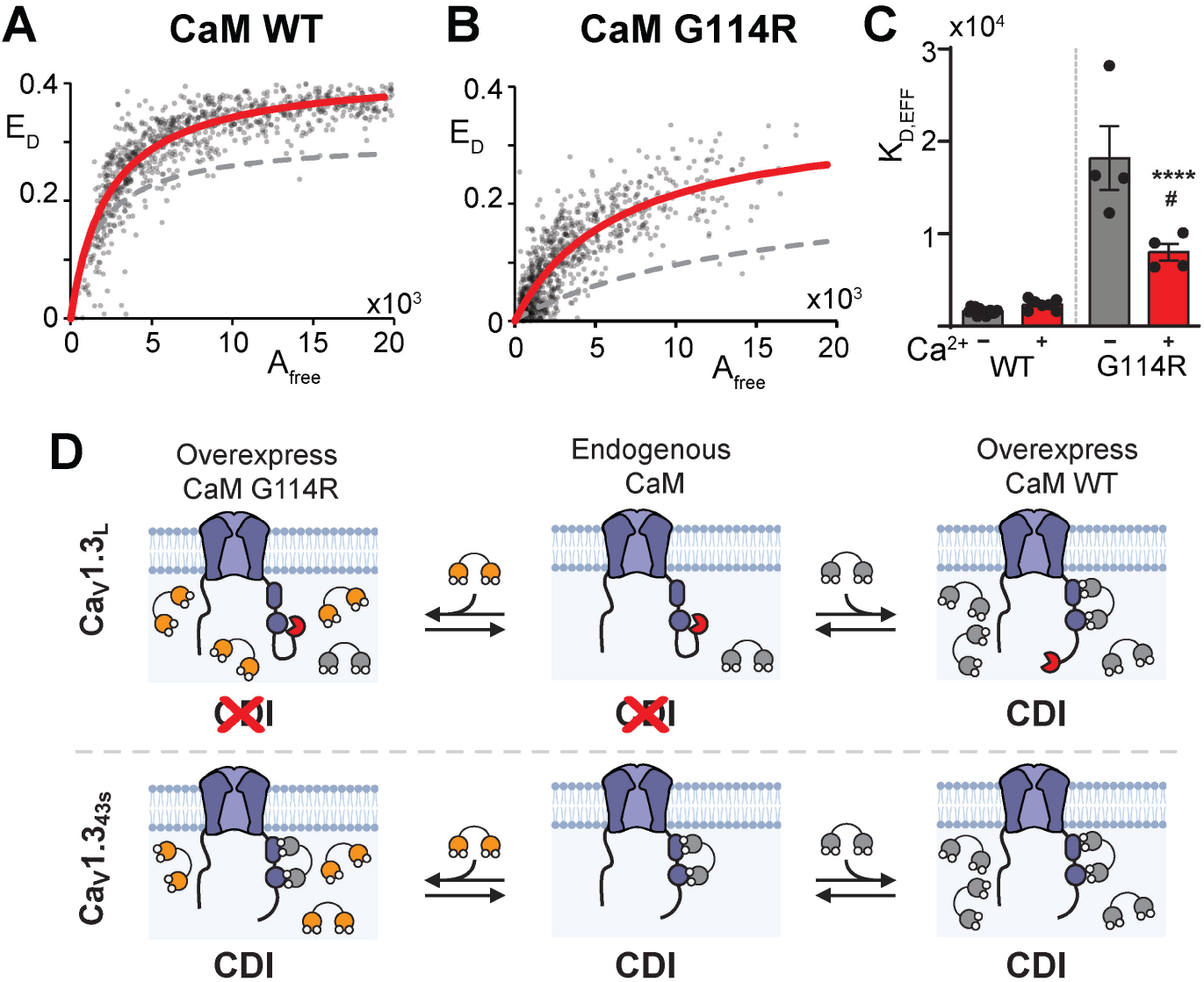
Binding of CaM G114R to Ca_V_1.3 is partially restored by elevated Ca^2+^. **A-B:** FRET binding curves from a live-cell flow cytometric FRET assay for the Ca_V_1.3 channel fragment and CaM WT (**A**) or CaM G114R (**B**) performed in high Ca^2+^ conditions. Each point reflects the result from an individual cell, with a curve in red fitted to estimate effective *K*_D_ (*K*_D,eff_). Low (basal) Ca^2+^ results are reproduced as the gray dashed curve for reference. **C**: Effective *K*_D_ values (*K*_D,eff_) plotted for the Ca_V_1.3 channel fragments, CaM WT, and CaM G114R under conditions of minimal (grey) or elevated cytosolic Ca^2+^ (red). (****, p > 0.0001 compared to WT in high Ca^2+^; #, p = 0.029 compared to apoCaM G114R). Data plotted ± SEM. **D**: Proposed mechanism under which endogenous WT CaM (grey) is capable of binding to Ca_V_1.3 and conferring CDI in the presence or absence of nonbinding CaM G114R (orange). In Ca_V_1.3_L_ with the ICDI/CTM (red) present, CDI occurs only when CaM WT, but not CaM G114R is overexpressed. In contrast, CaM WT, either endogenous or overexpressed, confers CDI in the absence of ICDI/CTM in Ca_V_1.3_43S_.

In addition to investigating the interaction between CaM and the Ca_V_1.3 IQ in low and high Ca^2+^ conditions, we explored the Ca^2+^-dependency of the interaction of each CaM variant with the channel domain. This was done by monitoring the change in fluorescence anisotropy of the 5-TAMRA-labeled Ca_V_1.3 IQ during titration with CaM at eight different Ca^2+^-concentrations (**Fig 7**), as previously done for the Ca_V_1.2 IQ (Wang 2018; Wang 2020; Brohus 2021). The affinity of CaM WT for Ca_V_1.3 IQ dramatically increased as a function of Ca^2+^-concentration (**Fig 7A**). Likewise, the affinity of all other CaM variants increased with Ca^2+^-concentration (**Fig 7B-G**). As expected, the Ca^2+^-insensitive variant, CaM_1234_, retained an apo-affinity for the Ca_V_1.3 IQ across the entire range of Ca^2+^ concentrations tested, demonstrating its inability to bind Ca^2+^-ions (**Fig 7H**). To summarize the effects of the CaM mutations on the interaction with the Ca_V_1.3 IQ, the apo-affinity, the Ca^2+^-affinity, and the Ca^2+^-sensitivity of each complex were determined (**Fig 7I-K**). In support of the FRET experiments, and the hypothesis of reduced channel preassociation, CaM G114R had the largest effect on the apo-affinity, causing a significant 10-fold reduction compared to CaM WT (**Fig 7I**). CaM D130G and D132G significantly reduced the apo-affinity by 2-fold compared to CaM WT. Of note, only CaM F142L caused an increase in the apo-affinity (13-fold compared to CaM WT), in agreement with what has been observed for the Ca_V_1.2 IQ (Wang et al., 2018). In Ca^2+^ saturating conditions, CaM D130G reduced the affinity of the complex 5.5-fold compared to CaM WT whereas CaM D132G and F142L caused a 3.5- and 2-fold increase, respectively (**Fig 7J**). As a measure of the Ca^2+^-sensitivity of CaM when in complex with the Ca_V_1.3 IQ domain, we determined the Ca^2+^-concentration causing a half-maximal increase in complex affinity (EC50) (**Fig 7K**). The EC_50_-value for CaM WT/Ca_V_1.3 IQ is 400 nM free Ca^2+^. All C-lobe mutations impaired the Ca^2+^-sensing of CaM when complexed to the Ca_V_1.3 IQ (2-11-fold) in line with their reduction of C-lobe Ca^2+^-affinity (**Fig 7K**), and consistent with their effect in impairment of Ca_V_1.3 CDI.

**Figure 7:**
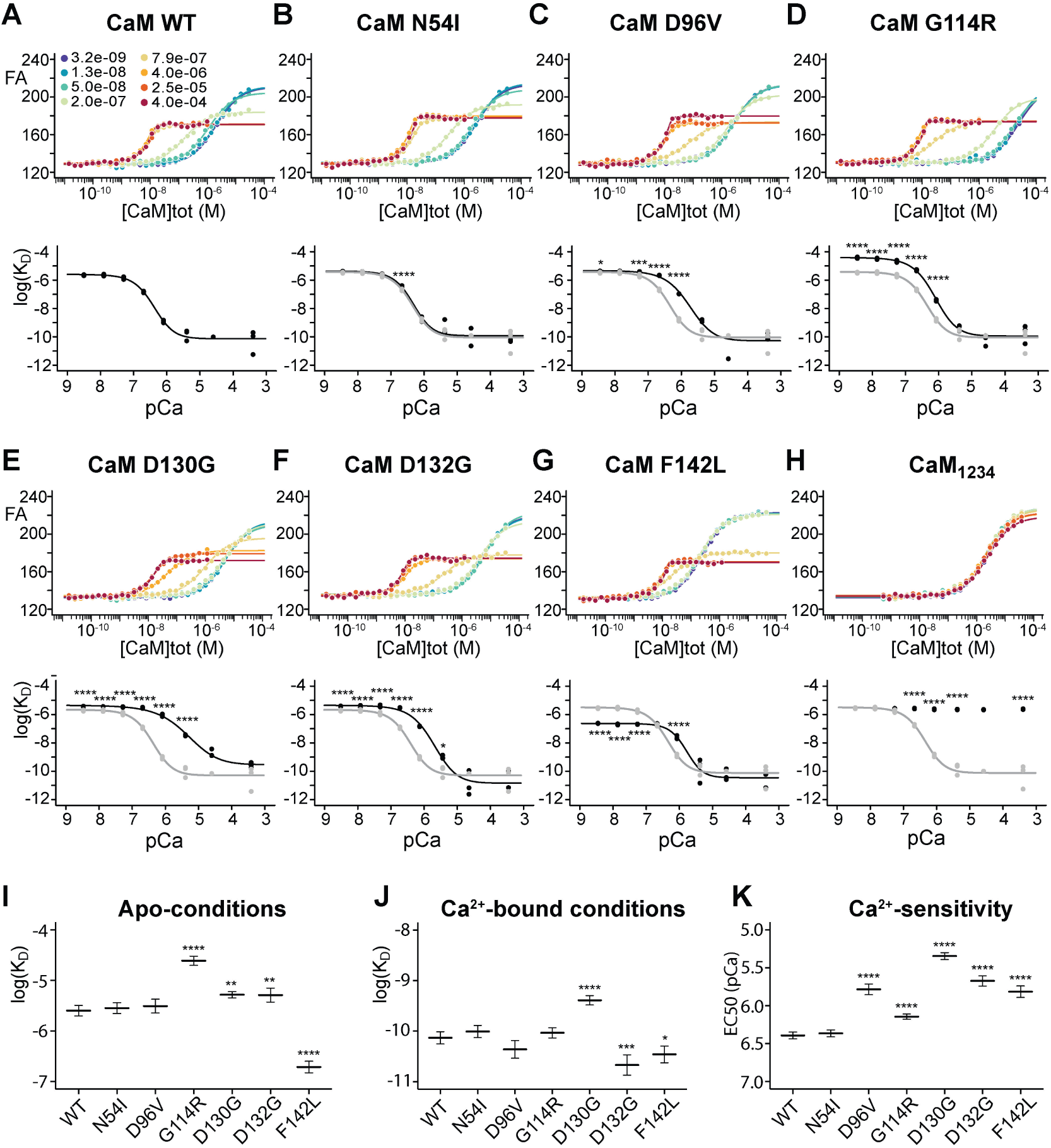
Calmodulin mutations impair the Ca^2+^ sensitivity of CaM in complex with the Ca_V_1.3 IQ domain. **A-H**, Top: exemplar experiment for the variant in question showing the development in fluorescence anisotropy of the TAMRA-labeled Ca_V_1.3 IQ-peptide as a function of CaM concentration. A stoichiometric binding model was fitted to the data to produce a binding curve and to determine the affinity of the interaction. Different colors represent different free Ca^2+^ concentrations, ranging from 3 nM to 0.4 mM. Bottom: The CaM/IQ -peptide affinity (*K*_D_) as a function of free Ca^2+^ concentration (shown as -log10([Ca^2+^]_free_)); the variant in question is shown in black with the WT profile shown in grey. **I-J**, comparison of the Ca_V_1.3 IQ affinity of all CaM variants at apo- and Ca^2+^-saturating conditions, respectively (shown as log10(*K*_D_)). **K**, Calmodulin variant Ca^2+^ sensitivity in the presence of IQ peptide, determined as the EC50 Ca^2+^ concentration (fitted to the change in log10(*K*_D_)). Note that for CaM_1234_, which cannot bind Ca^2+^-ions, no EC50 curve was fitted.

### The Impact of Pathological CaM mutations on Ca_V_2.1 Regulation

Having evaluated the impacts of pathological CaM mutants on Ca_V_1.3, we next turned to investigate the P/Q-type channel, Ca_V_2.1. To do so, we utilized a similar experimental setup as before with Ca_V_2.1 transfected into HEK293 cells along with the mutant CaMs. For Ca_V_2.1, C-lobe mediated CaM regulation takes the form of CDF (DeMaria et al., 2001). To evaluate CDF, we employed a step depolarization, with and without a prepulse at varying potentials. Ca^2+^ entry during the prepulse initiates CDF, resulting in a larger initial current during the test pulse. To quantify the amount of CDF, the area between the two curves is integrated to give the relative facilitation (RF) value (**Fig. 8**). As expected for a C-lobe mediated process, the N-lobe CaM mutant N54I did not significantly impair Ca_V_2.1 CDF as compared to WT (**Fig. 8B**). On the other hand, the C-lobe mutants G114R, D130G, D132G, and F142L all significantly impaired CDF, with G114R and F142L displaying somewhat more moderate effects (**Fig. 8D-G**). The artificial mutant CaM_1234_ demonstrates the impact of a complete loss of Ca^2+^ binding to CaM (**Fig 8H**). Interestingly, however, the C-lobe mutant D96V did not display impaired CDF (**Fig. 8C**) despite the fact that the mutated residue resides in the third Ca^2+^ binding EF hand and has been shown to dramatically reduce Ca^2+^ binding (Limpitikul et al., 2014; Sondergaard et al., 2015b). We therefore wondered if the affinity of the D96V mutant for Ca_V_2.1 might be impaired as seen for G114R for Ca_V_1.3.

**Figure 8:**
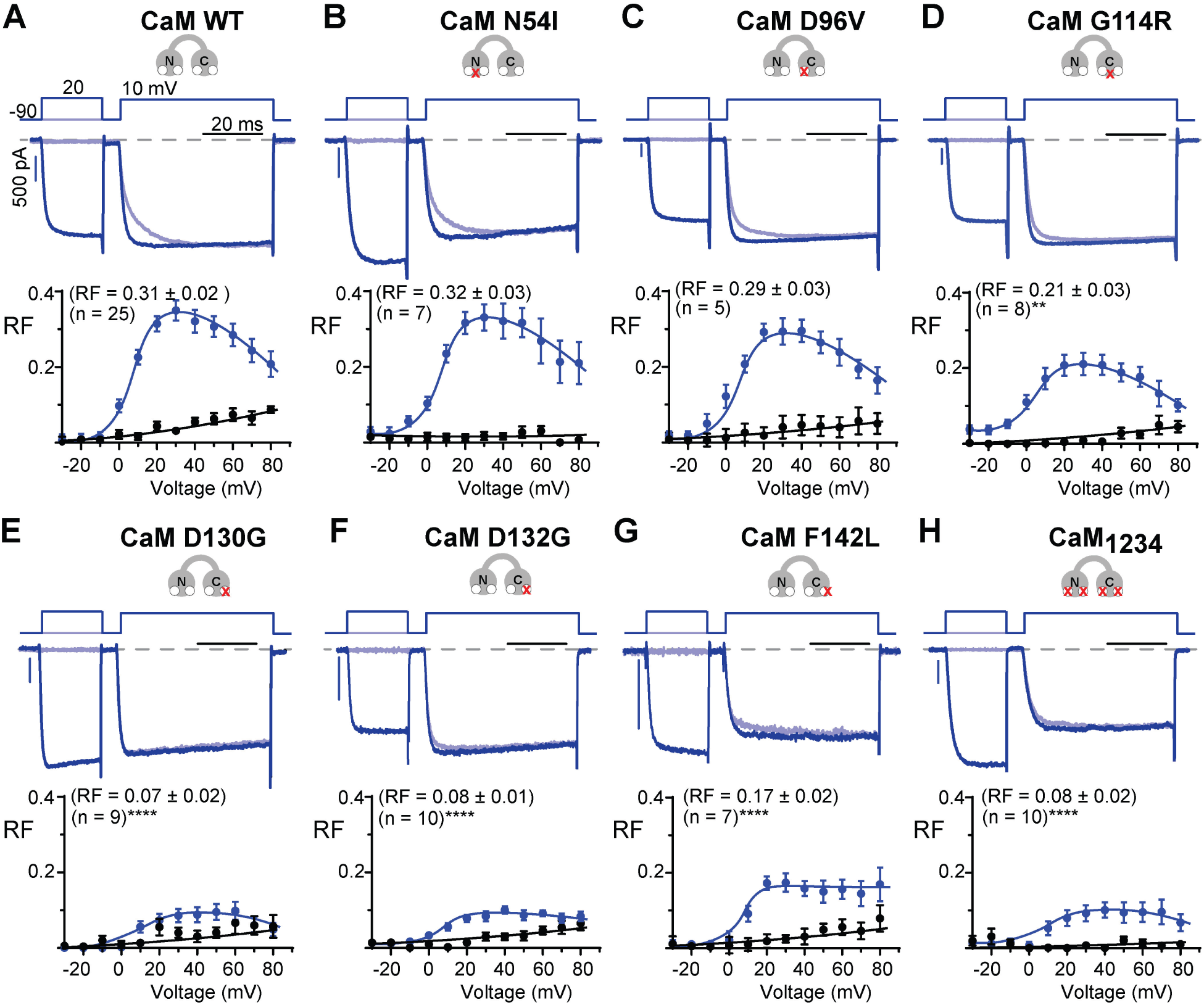
Calmodulin mutants impair Ca_V_2.1 CDF **A-H**, Top: Exemplar current traces for Ca_V_2.1 co-transfected with the indicated CaM variant. Test depolarization was performed with (dark blue) or without (light blue) a prepulse to 20 mV. Ca^2+^ currents recorded without a prepulse are normalized to Ca^2+^ currents with a prepulse, with scale bars referring to prepulse currents. Bottom: data displaying RF values for Ca_V_2.1 in Ca^2+^ (blue) and Ba^2+^ (black) and the indicated CaM variants recorded at varying test potential voltages (blue). RF values were compared at 10 mV for each of the CaM variants using a one-way ANOVA and Dunnett’s multiple comparison test relative to CaM WT (****, *p* < 0.0001). Data plotted ± SEM.

As before, we evaluated the capacity of mutant CaMs to bind via FRET. Using a C-terminal fragment of Ca_V_2.1 coupled to a cerulean fluorophore, the donor molecule of the FRET pair was constructed. Similarly, each mutant CaM was ligated to Venus as the acceptor fluorophore. Evaluation of the FRET binding curves to estimate *K*_D,eff_ under Ca^2+^ free conditions revealed that CaM G114R alone displayed significantly altered binding affinity (**Fig. 9A-G**), although the effect was much milder as compared to the binding deficits observed in Ca_V_1.3. Interestingly, CaM_1234_ displayed a similar mild but significant increase in *K*_D,eff_ (**Fig. 9J**), implying that even in the apo state, this form of CaM has diminished binding affinity.

**Figure 9:**
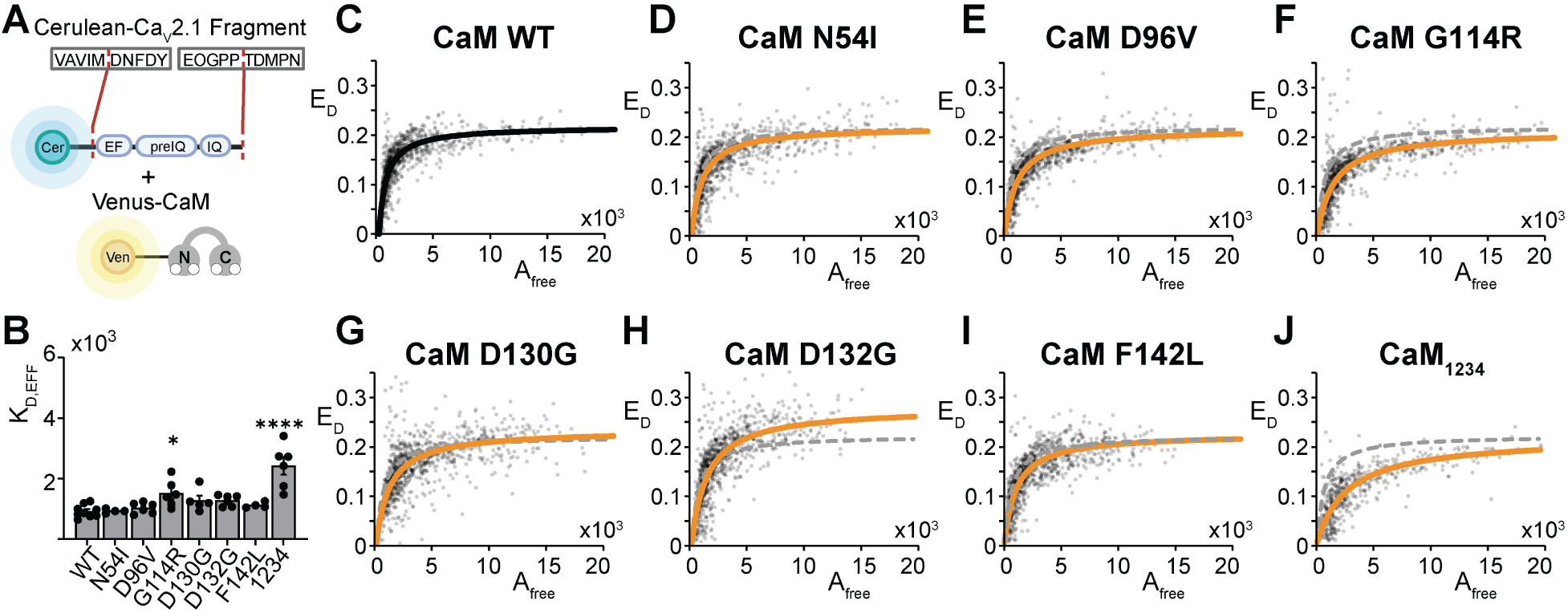
Binding of CaM to Ca_V_2.1. **A:** A Ca_V_2.1. C-terminal channel fragment including the IQ domain as indicated is concatenated with Cerulean. Each CaM variant was generated within a Venus-CaM construct. **B**: Measured relative binding efficiencies (*K*_D,eff_) plotted for the Ca_V_2.1 channel fragment and CaM variants in low cytosolic Ca^2+^. Significance was ascertained with a one-way ANOVA followed by a Dunnet’s test for multiple comparisons relative to WT (****, *p* > 0.0001) (*, *p* = 0.0142). Data plotted ± SEM. **C-J**: FRET binding curves from a live-cell flow cytometric FRET assay for the Ca_V_2.1 channel fragment and indicated CaM variant. Each point reflects the result from an individual cell, with a curve in black (WT) or orange (mutant) fitted to estimate effective *K*_D_ (*K*_D,eff_). Data is plotted as a function of free acceptor fluorophore molecules (*A_f_*_ree_) and donor-centric FRET efficiency (*E*_D_). WT results are reproduced on variant data as the gray dashed curve for comparison.

To explore the Ca^2+^-dependency of the CaM/Ca_V_2.1 IQ interaction, we once again used the fluorescence anisotropy assay (**Fig 10**). The affinity of CaM WT for the Ca_V_2.1 IQ dramatically increased with increasing Ca^2+^-concentration (**Fig 10A**). All other CaM variants, except CaM_1234_, followed the same pattern of increasing complex affinity with Ca^2+^-concentration (**Fig 10B-H**). In the low Ca^2+^ apo-state, all CaM C-lobe mutations displayed a small but significant reduction in binding affinity for the Ca_V_2.1 IQ domain (1.2- to 2.7-fold compared to CaM WT) (**Fig 10I**). Of note, the magnitude of reduction in apo-affinity for CaM G114R towards the Ca_V_2.1 IQ was less than that towards the Ca_V_1.3 IQ (2.7-fold reduction for Ca_V_2.1 IQ vs 10-fold reduction for Ca_V_1.3 IQ). This difference in binding affinity between the two channels supports a lack of apoCaM G114R preassociation with Ca_V_1.3, and thus a lack of effect on CDI, while retaining an effect on Ca_V_2.1 CDF. Similarly to the observation for the apo-affinities, all C-lobe mutations caused a reduction in the affinity of the Ca^2+^-saturated complex (2- to 14-fold compared to CaM WT) (**Fig 10J**). All C-lobe mutations reduced the Ca^2+^-sensitivity of the complex (1.8- to 4.3-fold compared to CaM WT), in line with their reduction of C-lobe Ca^2+^-affinity (**Fig 7K**), and consistent with their general impairment of Ca_V_2.1 CDF. These results, however, do not explain the distinct effects of D96V observed in functional assays. It is possible that the relatively small Ca_V_2.1 peptide used here and in our FRET assay is not sufficient to elucidate all CaM binding effects.

**Figure 10:**
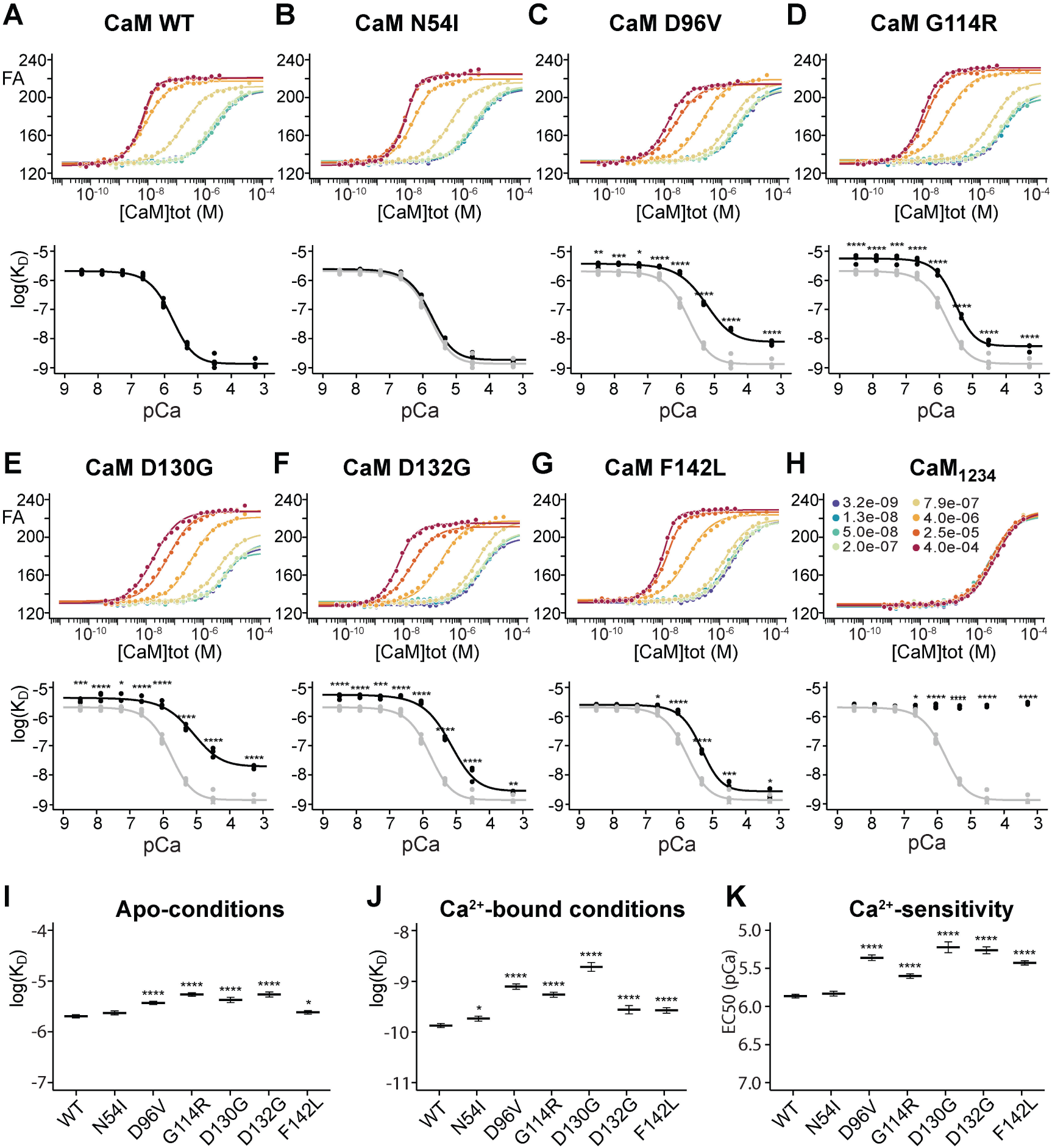
Calmodulin mutations impair the Ca^2+^ sensitivity of CaM in complex with the Ca_V_2.1 IQ domain. **A-H**, Top: exemplar experiment for the variant in question showing the development in fluorescence anisotropy of the TAMRA-labeled Ca_V_2.1 IQ-peptide as a function of CaM concentration. A stoichiometric binding model was fitted to the data to produce a binding curve and to determine the affinity of the interaction. Different colors represent different free Ca^2+^ concentrations, ranging from 3 nM to 0.4 mM. Bottom: The CaM/IQ -peptide affinity (*K*_D_) as a function of free Ca^2+^ concentration (shown as -log10([Ca^2+^]_free_)); the variant in question is shown in black with the WT profile shown in grey. **I-J**, comparison of the Ca_V_2.1 IQ affinity of all CaM variants at apo- and Ca^2+^-saturating conditions, respectively (shown as log10(*K*_D_)). **K**, Calmodulin variant Ca^2+^ sensitivity in the presence of IQ peptide, determined as the EC50 Ca^2+^ concentration (fitted to the change in log_10_(*K*_D_)). Note that for CaM_1234_, which cannot bind Ca^2+^-ions, no EC50 curve was fitted.

## Discussion

Work in both heterologous systems and model organisms has demonstrated that many CaM mutations can impair Ca_V_1.2 channel regulation leading to LQTS. Much has been done to characterize the mechanisms by which these deficiencies in Ca_V_1.2 regulation can lead to life-threatening cardiac outcomes; however, Ca_V_1.2 channels, and indeed other Ca_V_ channel subtypes, are also widely expressed throughout the brain. Moreover, many calmodulinopathy patients show profound neurological symptoms. Importantly, the preassociation with apoCaM that was demonstrated to be important for Ca_V_1.2 (Limpitikul et al., 2014) is shared by other VGCCs, making them particularly vulnerable to CaM mutations which alter Ca^2+^ binding. Thus, multiple VGCCs may be implicated in the neurological pathogenesis of calmodulinopathies.

The precise regulation of VGCCs in neurons is known to be critical for the physiological functions of the nervous system. Impairment of Ca_V_1.3 inactivation is observed in multiple neurological disorders including autism spectrum disorder (Limpitikul et al., 2016; Pinggera et al., 2018) and epilepsy (Pinggera et al., 2017). We find that a number of previously identified CaM mutants impair the CDI of these channels across distinct splice backgrounds. In particular, we find CaM mutants decrease CDI in Ca_V_1.3_L_, which possess the modulatory ICDI/CTM domain and is expressed abundantly in the heart and brain. Likewise, these CaM variants impair CDI in Ca_V_1.3_43s_, a variant lacking ICDI/CTM and expressed at higher levels in the brain as compared to the heart (Bock et al., 2011). Similarly, Ca_V_2.1 channels are widely expressed in neurons (Bourinet et al., 1999; Pietrobon, 2010) and contribute to short-term synaptic plasticity and rhythmicity (Catterall et al., 2013; Nanou et al., 2018). Disruption of these channels leads to severe symptoms, including epilepsy, migraine and neurodevelopmental delay (Pietrobon, 2010). Here, we show that many CaM mutations can impair the regulation of multiple VGCCs, including Ca_V_1.2, Ca_V_1.3, and Ca_V_2.1.

The CaM N54I mutation has a well-evidenced pathological role in CPVT (Nyegaard et al., 2012) with the effects largely independent of VGCC channels, making it a useful control for nonspecific effects. Our binding (FRET and fluorescence anisotropy) data demonstrates that it is able to readily associate with Ca_V_1.3 and Ca_V_2.1. However, it fails to impair the CDI of Ca_V_1.3 or the CDF of Ca_V_2.1. Interestingly, Ca_V_1.3 channels exhibit both N- and C-lobe mediated CDI, thus the N-lobe locus of N54I does not exclude a strong effect on the channel. However, the preservation of CDI and CDF is consistent with prior data showing that N54I has a milder impact on CaM binding to Ca^2+^ than LQTS-associated mutations shown to act on Ca_V_1.2 (Sondergaard et al., 2015a, Vassilakopoulou, 2015 #1679). Moreover, this lack of VGCC effects aligns with the lack of documented neurological symptoms in patients harboring this mutation (Crotti et al., 2023).

CaM mutations at residue G114 have previously been associated with sudden death and cardiac arrest, however, neurological symptoms were noted to be absent (Brohus et al., 2021; Crotti et al., 2023). G114R resides in the C-lobe of CaM and has been shown to impair Ca^2+^ binding in a similar fashion to other pathogenic mutations linked to cardiac arrhythmias, although with less not at severity as compared to D96V, D130G, D132G or F142L (Brohus et al., 2021). CaM G114R has also been shown to impair Ca_V_1.2 CDI, although to a lesser extent as compared to the LQTS-associated CaM mutants described in this study (Brohus et al., 2021). While these facts might predict a similar impairment of Ca_V_1.3, our data suggest a different result. In Ca_V_1.3_L_, CaM G114R appears to diminish CDI and increase VDI as compared to overexpressed CaM WT (**Fig. 2**). However, in Ca_V_1.3_43s_, the mutation has no effect on CDI or VDI (**Fig. 4**). In addition, our FRET results indicate that CaM G114R does not bind Ca_V_1.3 effectively in the apo state, a result supported by fluorescence anisotropy data which specifically showed a prominent reduction of binding to the Ca_V_1.3 IQ for G114R at low Ca^2+^ concentrations. Thus, we postulate that CaM G114R does not preassociate with either Ca_V_1.3 splice variant at low Ca^2+^ and the results reflect the lower expression of endogenous CaM as compared to our overexpressed CaM WT control. Accordingly, the apparent CDI and VDI changes seen in Ca_V_1.3_L_ can be explained by a simple mass-action effect (**Fig. 7E**). Ca_V_1.3_L_ harbors an ICDI/CTM region known to compete with CaM for binding to the channel IQ, thus reducing CDI (Liu et al., 2010). Overexpression of CaM WT outcompetes ICDI, enhancing CDI in our control recordings. Comparing the CDI of Ca_V_1.3_L_ coexpressed with CaM G114R to prior recordings of this channel (Singh et al., 2008; Liu et al., 2010) show a profile nearly identical to that achieved with endogenous CaM levels. Importantly, this mechanism also explains the rapid CDI kinetics of the channel with CaM G114R, and the VDI effect. With both lobes of endogenous CaM available to initiate CDI, the kinetics reflect both the rapid C-lobe mediated CDI combined with the slower N-lobe component (Yang et al., 2002). Moreover, the presence of ICDI/CTM has been shown to enhance channel VDI, explaining the increase in VDI observed in the presence of CaM G114R as the effect of ICDI/CTM binding (Singh et al., 2008; Bock et al., 2011). Similarly, the WT levels of CDI seen in Ca_V_1.3_43s,_ which lacks the ICDI/CTM domain, are consistent with the function of endogenous WT CaM (Singh et al., 2008) in the presence of unbound CaM G114R.

Interestingly, CaM G114R does impair Ca_V_2.1 CDF under our experimental conditions of heterologous overexpression, although to a lesser extent as compared to other C-lobe mutants (**Fig. 8D**). This result is consistent with the modest reduction in binding affinity with the channel IQ (**Figs. 8, 9**), as well as the more moderate effect of this mutation on Ca^2+^ binding (Brohus et al., 2021). This is also reflected in a more moderate effect on the CaM Ca^2+^ sensing in the presence of the Ca_V_2.1 IQ domain, as shown by our fluorescence anisotropy experiments (**Fig 10K**). Thus, CaM G114R may not preassociate to the same extent as CaM WT, and CaM G114R retains some Ca^2+^ sensitivity while bound to the channel, enabling some CDF, compared to CaM D130G and D132G (**Fig 8D-G**). Thus, our data indicate that CaM G114R likely has minimal impacts on Ca_V_1.3 channel regulation and a modest potential to impair Ca_V_2.1 CDF, consistent with the lack of neurological symptoms described in patients. Moreover, these results highlight the differential impact a single CaM mutant can have on Ca_V_2.1 vs Ca_V_1.3.

D96V has been shown to severely impair the ability of CaM to bind Ca^2+^ (Vassilakopoulou et al., 2015) and has a significant impact on CDI in Ca_V_1.2 (Limpitikul et al., 2014) (**Fig. 1**) and Ca_V_1.3 (**Figs. 2, 4**). Although it binds well with both Ca_V_1.3 and Ca_V_2.1 channel fragments in our FRET assay, it fails to impair Ca_V_2.1 CDF (**Fig 7**). The unusual discrepancy between Ca_V_1.3 and Ca_V_2.1 channel regulation by CaM D96V is not made clearer by our fluorescence anisotropy results, where it shows a similar profile of Ca^2+^-dependent binding to a Ca_V_2.1 channel peptide as the other mutants. It is therefore tempting to speculate that this mutation may have an effect on structure that imparts a distinct effect in the context of Ca_V_1.2/1.3 vs. Ca_V_2.1. While such an effect remains to be elucidated, D96V has a documented effect on CaM structure, resulting in an unusual protein conformation that resembles neither apoCaM nor the fully calcified form of the protein (Wang et al., 2018). Interestingly, the structural changes of this mutation appear to be less severe as compared to those of the D130G mutation (Wang et al., 2020), indicating that perhaps Ca_V_2.1 CDF is more amenable to certain structural changes as compared to the Ca_V_1.2 or Ca_V_1.3 channels. It is also interesting to note, that while this mutation has a reported association with developmental delay (Crotti et al., 2013), the neurological effects appear less consistently reported (Crotti et al., 2013; Crotti et al., 2023) and were reported subsequent to cardiac arrest, making it difficult to rule out hypoxic injury. Due to the limited number of patients, it is difficult to assess whether the lack of Ca_V_2.1 effect by CaM D96V correlates with a reduction in neurological consequences, but it is possible that each of the channels (Ca_V_1.2, Ca_V_1.3 and Ca_V_2.1) may contribute in varying degrees to distinct neurological features.

The CaM mutations D130G and D132G follow a similar and predicted pattern: both are located within or just adjacent to an EF-hand in the C-lobe of CaM and severely impair the Ca^2+^-dependent regulation of Ca_V_1.3 and Ca_V_2.1. F142L, on the other hand, resides outside the EF-hand coordinating residues, but is critical to the packing and folding of the C-lobe of CaM. Interestingly, this mutation exhibits a comparable impact on CDI in the context of Ca_V_1.3 but exhibits a lesser effect on Ca_V_2.1 CDF as compared to the EF-hand mutations. Thus, the detailed mechanism underlying the loss of CaM regulation may be distinct for F142F. In addition, all three of these mutant CaMs show strong affinity for Ca_V_1.3 and Ca_V_2.1 channel peptides at low Ca^2+^ levels, indicating preservation of the preassociation feature. However, F142L again differs in that binding to Ca_V_1.3 (**Figs. 7,8**) and Ca_V_1.2 (Limpitikul et al., 2014; Wang et al., 2018) IQ region is enhanced at low Ca^2+^ levels (Crotti et al., 2013; Crotti et al., 2023). Each of these mutations have been documented in patients suffering from autism spectrum disorder, developmental delay, and/or epilepsy, with F142L associated with a broader ser of reported neurological deficits (Crotti et al., 2013; Kaplanis et al., 2020; Crotti et al., 2023). Thus, it is tempting to speculate that the enhanced low Ca^2+^ binding of this mutant to Ca_V_1.2 and Ca_V_1.3 may enable F142L to impact a greater number of Ca_V_1 channels even in the context of physiological expression in one out of 6 alleles, and this increased effect on this channel subtype may outweigh the lesser impact on Ca_V_2.1.

Neurological phenotypes differ widely among calmodulinopathy patients. This variability cannot be explained entirely by distinct mutation effects. The presence of three independent *CALM* genes encoding the protein leaves open the possibility of variable effects due to the fact that these genes are not expressed equivalently throughout different brain regions, tissues, and cell types. Moreover, the distinct impact on different VGCCs will be further modulated by neuronal context and channel expression, resulting in a complex combination of loss and gain of function effects. Additionally, while VGCCs represent an important pathogenic target, CaM is likely to modulate numerous targets, adding to the complexity of the pathogenesis. Thus, variability in neurological effects depends on multiple mechanisms and are thus difficult to predict. Finally, while deficits in Ca_V_ channel regulation are frequently characterized as “gain-of-function” (GoF) or “loss-of-function” (LoF) in terms of their impact on Ca^2+^ influx, the deficits imparted by the CaM mutations we have characterized are not so readily delineated into one of these categories. The same CaM mutation that impairs Ca_V_1.3 CDI, therefore permitting excess Ca^2+^ influx in what could be termed a GoF, can also impair Ca_V_2.1 CDF, resulting in diminished cytosolic Ca^2+^ and therefore a LoF. Due to these divergent impacts on two prevalent CaM targets, the effect of CaM mutations on overall neuronal Ca^2+^ signaling is likely to be multifaceted.

Overall, this study presents strong evidence that mutations in CaM can impact multiple VGCCs, expanding our knowledge of the potential pathogenic mechanisms underlying calmodulinopathies. The impact of CaM mutations on VGCC function incorporates many of the same principles identified in the disruption of Ca_V_1.2 CDI leading to severe LQTS, where VGCCs likely represent particularly vulnerable targets due to preassociation with CaM at low Ca^2+^ levels combined with a diminished response to increasing Ca^2+^-concentrations. Overall, disruption of VGCCs in the brain may represent an important component of the molecular mechanism for calmodulinopathies with neuronal impairment.

## Methods

### Transfection and molecular biology for electrophysiology

For electrophysiological experiments, a plasmid encoding a human α_1C_ channel variant containing exon 8a was employed for Ca_V_1.2 (Z34810) (Dick et al., 2008). Plasmids encoding a human α_1D_ channel variant containing exons 8a and 42 was employed for Ca_V_1.3_L_ (NG_032999) and a variant terminating in exon 43 was employed for Ca_V_1.3 43s (NM_001128839) (Sang et al., 2021) were previously provided by Dr. Jörg Striessnig (University of Innsbruck). A plasmid encoding an α_1A_ channel variant including exons 43 and 44 and excluding exon 47, originally provided by Dr. Terrance Snutch (University of British Columbia), was employed for Ca_V_2.1 (NM_001127221) (Lee et al., 2015). Each of these was within a pcDNA3 expression vector. For these same experiments, human *CALM1* cDNA within the pIRES2-EGFP vector was utilized for WT CaM (Clontech Laboratories). From this WT CaM-pIRES-EGFP construct, pathogenic CaM mutations were generated using QuikChange Lightning™ site-directed mutagenesis (Agilent). 8 μg each of human α_1_ cDNA, rat brain β_2a_ (M_80545) or rat brain β_1b_ (X61394), rat brain α_2δ_ (NM_012919.3) subunits, and WT or mutant CaM were heterologously co-expressed. For all recordings, β_2a_ was used unless otherwise specified. Expression was enhanced with 2 μg of simian virus 40 T antigen (Tag) cDNA. Expression of all plasmids was under the control of a cytomegalovirus (CMV) promoter. HEK293 cells were cultured in 10-cm dishes on glass coverslips, and channels were transfected using an established Ca^2+^ phosphate method (Peterson et al., 1999).

### Patch clamp recordings

Whole-cell voltage-clamp recordings of HEK 293 cells were carried out 1-2 days post-transfection at ambient temperature. An Axopatch 200B amplifier (Axon Instruments) was utilized to generate the recordings. Noise in these recordings was minimized via a lowpass filter at 2 kHz, followed by digital sampling at 10 kHz. Series resistances of 1.5-5 MΩ and well as P/8 leak subtraction were considered as necessary conditions. Internal solutions contained (in mM): CsMeSO3, 114; CsCl, 5; MgCl2, 1; MgATP, 4; HEPES (pH 7.4), 10; and BAPTA, 10; at 295 mOsm adjusted with CsMeSO_3_. External solutions contained (in mM): TEA-MeSO3, 140; HEPES (pH 7.4), 10; and CaCl2 or BaCl_2_, 40; at 300 mOsm, adjusted with TEA-MeSO3. These solutions produced the following uncorrected junction potentials: 10 BAPTA/40 Ca^2+^: 10.5 mV; 10 BAPTA/40 Ba^2+^: 10.2 mV. Note that the use of 10 BAPTA limits Ca^2+^ to only the local signal due to entry through the channel (Dick et al., 2008). For Ca_V_1.3 both lobes of CaM are able to respond to this local Ca^2+^ signal, but for Ca_V_2.1 this manipulation restricts CaM regulation to C-lobe mediated CDF (Ben-Johny and Dick, 2022), reducing the confounding effect of N-lobe mediated CDI in our CDF measurements. However, it should be noted that this manipulation also reduces the ability to observe n lobe effects, thus the N54 mutation is used largely as a negative control under these conditions.

### Quantification and statistical analysis of patch clamp data

For *r*_300_ Ca^2+^ data in Ca_V_1.2/CaM experiments, a ratio of peak currents during a step depolarization to 30 mV to the current remaining following 300 ms was employed for each CaM variant and WT. For each cell, this value was subtracted from the analogous value for Ba^2+^ traces at 30 mV, resulting in an *f*_300_ value that quantified CDI in Ca_V_1.2. The *f*_300_ value from each cell was averaged for the D132G mutant condition and were compared to the WT via Student’s t-test. For both sets of Ca_V_1.3/CaM datasets, similar comparisons were employed, with the following differences: currents were compared at 10 mV step depolarizations, a 1000 ms depolarization was used resulting in *f*_1000_ values. For experiments investigating Ca_V_1.3/CaM VDI, *r*_1000_ values acquired during Ba^2+^ depolarizations were used to estimate VDI.

For Ca_V_2.1/CaM experiments, CDF was characterized by comparing Ca^2+^ currents during depolarizing voltage steps to 10 mV in the presence and absence of a prepulse which was sufficient to partially facilitate the channel. (Chaudhuri et al., 2004). The area between the currents generated thus (Δ*Q*) divided by the time constant of the curve without the prepulse τ, gives an approximation of the degree of CDF triggered by the prepulse, termed relative facilitation (RF). In cases where the lack of CDF led to an absence of a monoexponential curve to derive τ from, τ was set as the average value from WT dataset (8.5). Data from prepulses to 20 mV were used for statistical comparisons. For all patch clamp experiments where more than one mutant was being compared to WT, the relevant values were compared via one-way ANOVAs using Dunnet’s multiple comparison corrections, with each mutant being compared to WT. Data from individual cells was excluded from analysis if peak Ca^2+^ currents were below 150 pA, or if the traces showed evidence of buffer depletion.

### Generation of FRET constructs

The principles underlying flow cytometric FRET having been previously established (Ben-Johny et al., 2016; Lee et al., 2016; Rivas et al., 2021). Venus-CaM plasmids correspond to those described previously (Ben-Johny et al., 2016; Lee et al., 2016; Niu et al., 2018). QuikChange Lightning mutagenesis (Aligent) was employed to create the Venus-CaM mutants of interest. For Ca_V_1.3, the C-terminal fragment (DNFDY…VGKYP) was ligated into a pcDNA3 expression vector containing an upstream Cerulean fluorophore. The DNA template encoding the Ca_V_1.3 fragment as well as XbaI and NotI restriction digest sites at positions flanking the coding sequence was purchased (Twist Bioscience). Using this template, digestion with these enzymes was carried out, followed by ligation into the expression vector using T3 DNA ligase (New England Biolabs). To ensure expression and read-through, the Cerulean fluorophore was preceded by a CMV promotor, and an in-frame short linker sequence of alanine was present between the end of the Cerulean sequence and before the Ca_V_1.3 fragment. At the end of the Ca_V_1.3 fragment was a stop codon to create the fusion protein suitable for FRET. The same procedure was employed to generate the Cerulean/Ca_V_2.1 construct, with the Ca_V_2.1 sequence (DNFEY…EOGPP) inserted into the vector. Dimers linking Cerulean and Venus of various lengths (C5V, C32V, C40V, CTV) were utilized as previously described (Ben-Johny et al., 2016; Lee et al., 2016; Rivas et al., 2021). All constructs were verified by sequencing (Plasmidsaurus).

### Transfection of FRET constructs

Cells were seeded in 6-well plates and transfected with FRET constructs and fluorophore pairs using a PEI protocol (Longo et al., 2013). 48 hours post-transfection, cells were treated with 100 μM cycloheximide for 2-4 hours to reduce incidence of immature fluorophores, and were then harvested and resuspended in Tyrode’s solution with minimal Ca^2+^. For experiments involving calcified CaM, 10 mM CaCl_2_ was added to equal volumes of cell suspension along with 4 μM ionomycin (Sigma-Aldrich I0634).

### FRET data collection and analysis

To estimate binding curves and *K*_D,eff_ for each of the donor-acceptor pairs, fluorescence measurements are acquired from three filters within the flow cytometer, specifically selected to detect donor, acceptor, and FRET signals. This technique, termed 3^3^-FRET, permits the isolation of FRET signal from donor and acceptor signals (Erickson et al., 2001). This methodology required evaluation of instrument-specific properties during each experimental run, evaluated using a series of linked Cerulean/Venus dimers such that a 1:1 stoichiometry of each fluorophore was ensured (Ben-Johny et al., 2016). Moreover, Venus and Cerulean fluorophores, individually and coexpressed, were measured to permit evaluation of non-specific fluorescence that could confound FRET analysis.

Analysis of flow cytometric FRET was performed as previously described (Ben-Johny et al., 2016; Lee et al., 2016; Rivas et al., 2021). Net decreases in donor fluorescence due to FRET was computed from the signals from each of the specific filter cubes utilized. These decreases are analogous to FRET efficiencies acquired from acceptor photobleaching or donor quenching approaches, and correspond to the donor-specific FRET measurement, *E*_D_. For each of the thousands of cells within each sample, *E*_D_ was plotted against the concentration of unbound acceptor molecules (*A*_free_) in order to generate a binding curve, and effective *K*_D_ (*K*_D,eff_) was then calculated from this curve. *K*_D,eff_ values from each FRET experiment were compared using one-way ANOVAs using Dunnet’s multiple comparison corrections, with each mutant being compared to WT. Data were analyzed by custom MATLAB software (Ben Johny, 2020) (Mathworks, MA).

### Fluorescence Anisotropy Binding Assays

Peptides representative of the Ca_V_1.3 and Ca_V_2.1 IQ-domains were N-terminally labeled with a 5-TAMRA fluorophore (Ca_V_1.3: 5-TAMRA-DEVTVGKFYATFLIQDYFRKFKKRKEQGLVGKYP-NH2, Ca_V_2.1:_5-TAMRA-TDLTVGKIYAAMMIMEYYRQSKAKKLQAMREEQD-NH2; Proteogenix, purity > 95%). Using a previously described 384-well plate-based assay (Brohus et al., 2019), CaM was titrated across the 24 columns at 8 different free calcium concentrations (3 nM to 400 µM), one per row (Brohus et al., 2021). The change in fluorescence anisotropy of the TAMRA-labeled IQ-domains, a measure of their tumbling rate, was monitored as a function of CaM concentration. This setup resulted in 8 CaM/IQ-domain binding curves, each representing binding at a specific free Ca^2+^ concentration.

A stoichiometric binding model was fitted to each binding curve to determine the CaM/IQ-domain binding affinity (represented by the dissociation constant, *K*_D_) at each Ca^2+^ concentration (Jensen et al., 2024). In the case of Ca_V_1.3 IQ, binding curves for CaM G114R did not reach an upper plateau at low Ca^2+^-concentrations; thus, curve spans from CaM WT were assumed for these curves to determine the apo-affinity. Affinities and Ca^2+^ concentrations were log10-transformed before fitting an EC50 curve to determine the apo- and Ca^2+^-affinity of the CaM/Ca_V_ IQ interaction, and to determine the Ca^2+^-concentration that provides the half-maximal change in affinity between the apo- and Ca^2+^-bound states – a measure of the Ca^2+^-sensitivity of the interaction. In summary, the assay provides information on the Ca^2+^-dependency of the CaM/IQ-domain interaction across the physiologically relevant Ca^2+^-concentration span.

Curve fitting and data plotting was done using a custom-written script pipeline in R (v. R-4.2.1). Statistical testing was done on log10-transformed affinities and EC50-values in GraphPad Prism 6 (v. 6.07) using a one-way ANOVA (WT as reference) with Dunnett’s *post hoc* test for multiple comparisons. Significance levels were indicated by asterisks: **** (p-value < 0.0001), *** (p-value < 0.001), * (p-value < 0.05).

## Acknowledgements

We thank all members of Dr. Dick’s lab for helpful suggestions and discussion of the project. We also thank Josiah Owoyemi and Emma Gudmundsson for dedicated technical support. This work was supported by grants from the National institutes of Neurological Disorders and Stroke (R21NS127294 to IED) and the National Heart Lung and Blood Institute (R01HL149926 to IED and R01HL163576 to MBJ).

